# Absence of the Z-disc protein α-actinin-3 impairs the mechanical stability of *Actn3KO* mouse fast-twitch muscle fibres without altering their contractile properties or twitch kinetics

**DOI:** 10.1101/2021.11.09.467867

**Authors:** Michael Haug, Barbara Reischl, Stefanie Nübler, Leonit Kiriaev, Davi A.G. Mázala, Peter J. Houweling, Kathryn N. North, Oliver Friedrich, Stewart I. Head

## Abstract

**Background:** A common polymorphism (R577X) in the *ACTN3* gene results in complete absence of the Z-disc protein α-actinin-3 from fast-twitch muscle fibres in ~16% of the world’s population. This single gene polymorphism has been subject to strong positive selection pressure during recent human evolution. Previously, using an *Actn3KO* mouse model, we have shown in fast-twitch muscles, eccentric contractions at L_0_+ 20% stretch did not cause eccentric damage. In contrast, L_0_+ 30% stretch produced a significant ~40% deficit in maximum force; here we use isolated single fast-twitch skeletal muscle fibres from the *Actn3KO* mouse to investigate the mechanism underlying this.

**Methods:** Single fast-twitch fibres are separated from the intact muscle by a collagenase digest procedure. We use label-free *second harmonic generation* (SHG) imaging, ultra-fast video microscopy and skinned fibre measurements from our *MyoRobot* automated biomechatronics system to study the morphology, visco-elasticity, force production and mechanical strength of single fibres from the *Actn3KO* mouse. Data are presented as means ± SD and tested for significance using ANOVA.

**Results:** We show that the absence of α-actinin-3 does not affect the unloaded maximum speed of contraction, visco-elastic properties or myofibrillar force production. Eccentric contractions demonstrated that chemically skinned *Actn3KO* fibres are mechanically weaker being prone to breakage when eccentrically contracted. Furthermore, SHG images reveal disruptions in the myofibrillar alignment of *Actn3KO* fast-twitch fibres with an increase in Y-shaped myofibrillar lattice shifts.

**Conclusions:** Absence of α-actinin-3 from the Z-disc in fast-twitch fibres disrupts the organisation of the myofibrillar proteins, leading to structural weakness. This provides a mechanistic explanation for our earlier findings that, *in vitro* intact *Actn3KO* fast-twitch muscles are significantly damaged by L_0_+ 30%, but not, L_0_+ 20%, eccentric contraction strains. Our study also provides a possible mechanistic explanation as to why α-actinin-3 deficient humans have been reported to have a faster decline in muscle function with increasing age, that is; as sarcopenia reduces muscle mass and force output, the eccentric stress on the remaining functional α-actinin-3 deficient fibres will be increased, resulting in fibres breakages.

## Background

Around 16% of humans lack α-actinin-3, due to a homozygosity for a common polymorphism in the *ACTN3* gene. This single gene polymorphism has been subject to strong positive selection during the last 50,000-60,000 years corresponding to the migration of modern humans from the African continent (1, 2). Intriguingly, two recent publications suggest that a major positive selection pressure may have been the fact that the α-actinin-3 polymorphism improves an individual’s cold acclimatization (3, 4). The *ACTN3* gene has become known colloquially as the “gene for speed” (1, 5). α-Actinin-3 deficiency is not associated with any skeletal muscle pathology, indeed it appears to be beneficial for female elite endurance athletes (6), however, it should be noted several subsequent studies in humans (which may be underpowered due to the large genetic variability) have not supported this first report (7). The speed of shortening of a muscle fibre depends largely on the myosin heavy chain (MyHC) isoform present (8). In previous studies we have shown that MyHC expression is unaltered in *Actn3 knockout (KO)* fibres (9) and by using a skinned fibre preparation, demonstrated that there is no difference between *Actn3KO* and wild type (WT) fast-twitch fibres regarding the Ca^2+^ sensitivity of the contractile proteins (10). In an intact preparation, using a high-speed imaging technique (8) and enzymatically isolated single fibres from *Actn3KO* and WT mice, we showed no difference in maximum speed of unloaded shortening during a single action potential triggered twitch (3). Taken as a whole, these data suggest the *ACTN3KO* gene does not alter the myosin isoform or contractile functioning of the contractile proteins.

The α-actinins are rod-shaped proteins of 35nm length that form antiparallel homodimers. Mammalian skeletal muscle expresses α-actinin-2 and -3. These isoforms are the major component of the Z-disc. α-Actinin-2 comprises the Z-discs of slow-twitch muscles while α-actinin-3 is found exclusively in the Z-discs of fast-twitch muscles (11). The Z-discs play a key role in longitudinal force transmission from the sarcomeres to the tendons (5). In human fast-twitch muscles, α-actinin-3 is more abundant in type 2X fibres compared to type 2A (12). Fast-twitch fibres are particularly susceptible to damage from eccentric contractions while slow-twitch fibres are very resistant to any damage from eccentric contractions (13, 14). While it is not clear why fast-twitch fibres are more susceptible to damage due to eccentric contractions compared to slow-twitch fibres, one structural reason comes from the observations that fast-twitch fibres have narrower Z-discs which would provide less mechanical support, i.e. higher mechanical stress, during high tension contractions (11). The width of the Z-disc largely reflects the amounts of α-actinin proteins anchoring the actin filaments of adjacent sarcomeres at the Z-discs. In *ACTN3KO* muscles of XX individuals there is a complete absence of α-actinin-3 which is functionally compensated for by the closely related protein α-actinin-2 (1). If α-actinin-2 configures in the narrow Z-disc fast-twitch profile in XX individuals, then these Z-discs may be less stable than the “wild-type” narrow α-actinin-3 fast-twitch Z-discs present in RR individuals homozygous for the *ACTN3* gene. In our earlier studies (15, 16) on isolated intact fast-twitch extensor digitorum longus (EDL) muscles from our mouse *Actn3KO* models, we compared the effect of eccentric contractions at L_0_+20% and L_0_+30% stretch. Eccentric contractions at L_0_+20% stretch did not result in a significant eccentric damage force deficit, in contrast L_0_+30% stretch did produce a significant ~40% force deficit. We interpreted these results as suggestive that an absence of α-actinin-3 increases the susceptibility to damage when *Actn3KO* mouse fast-twitch muscles are subject to high forces. However, in intact muscles, there is an interference from intermuscular pathways of lateral force transmission via the dystrophin and desmin pathways as well as the mechanical role played by the connective tissue lattice supporting muscle fibres within the intact muscle. In the current study, we used single fibres to directly probe the effects of the absence of α-actinin-3 on the longitudinal mechanical strength and contractility of fast-twitch fibres.

## Methods

### Animals

The *Actn3*KO mouse line was previously created in this laboratory (17), and experiments were performed on male animals at 12-15 months of age. A total of 6KO and 6WT mice were used in the present study. Use of animals was approved by the Animal Care and Ethics Committees of the Children’s Medical Research Institute and the University of New South Wales.

### Skeletal Muscle single fibre enzymatic isolation

Flexor digitorum brevis (FDB) and extensor digitorum longus (EDL) muscles were digested in Krebs solution composed of (in mM): 4.75 KCl, 118 NaCl, 1.18 KH_2_PO_4_, 1.18 MgSO_4_, 24.8 NaHCO_3_, 2.5 CaCl_2_ and 10 glucose containing 3 mg/ml collagenase type IV A (Sigma Aldrich, USA), gently bubbled with carbogen (95% O_2_, 5% CO_2_) and maintained at 37°C. After 25-30 minutes muscles were removed from the digest solution with a wide bore glass pipette and serially rinsed twice in Krebs solution containing 0.1% foetal calf serum. Single fibres were dispersed by gentle trituration. The FDB fibres were maintained in Krebs solution with 0.1% foetal calf solution at room temperature 21-23°C and continuously bubbled with carbogen. Using a pipette, 0.5 ml of solution was drawn and placed on a cleaned glass slide on an inverted microscope, each 0.5 ml contained between 10-50 fibres. FDB fibres attached firmly to the glass cover slip and were continually superfused with Krebs bubbled with carbogen at a rate of around 0.5 ml per minute. The FDB fibres were visualized at 200x magnification on a Nikon Eclipse Ti2-E Inverted Research Microscope. For fibre length and diameter measurements (Supplementary figure A), a grid was placed in the eye piece of the microscope so that it occupied ~50% of the field of view and all fibres in this view were recorded and processed using ImageJ open-source software, the microscope was calibrated using a stage micrometre, and a total of 200 WT FDB fibres were measured. Post-digest EDL muscles were rinsed first in Krebs with 0.1% foetal calf serum to stop the collagenase reaction and then rinsed for a second time in Krebs with no foetal calf serum and no added calcium before being placed in a relaxing solution with the following composition (mM): 117 K^+^, 36 Na^+^, 1 Mg^2+^, 60 HEPES, 8 ATP, 50 EGTA (Note: as the fibres are effectively chemically skinned by the high EGTA concentration, this is an intracellular solution). Transfers between solutions were made by sucking the digested muscle mass into a wide bored pipette. Finally, the muscle was gently agitated using a wide bore pipette to release individual fibres from the muscle. Fibres were maintained in the relaxing solution at four degrees centigrade for up to four hours before use.

### High-speed acquisition of transillumination images

We selected FDB fibres with a width of 35 micrometres or greater (supplementary figure A), FDB is a fast-twitch muscle and we only used fibres which responded briskly and repeatedly to a 1msec activating pulse, over 90% of FDB fibres are fast-twitch, however, we occasionally came across fibres which were slower to contract and relax (visual inspection), these fibres were not used (18). Intact single FDB fibres were electrically field-stimulated with supramaximal voltage pulses of 1 ms duration, 10 V amplitude over a range of frequencies from 10 Hz to 100 Hz. The stimulator probe was bipolar, with two fine platinum wires isolated up to the ends, the wires were attached to a fine Perspex rod mounted on a micromanipulator to enable it to be placed close (~10μm) to the neuromuscular junction of the selected FDB fibre. A CMOS PCO1200hs high-speed camera (PCO AG, Kehlheim, Germany) was mounted to the camera side-port of the Nikon inverted microscope. The Peltier-cooled camera was connected to a computer for acquisition control and data storage. Single fibres approximately covered a 520×160 pixel area when visualised through a 20x objective which allowed frame rates for shortening sequences of 4,200 frames per second. Recordings were synchronised with the induction of a single twitch and image read-out and storage from the ring-buffer of the camera was performed offline. For offline analysis of each experiment, an image sequence of approximately 1,000 to 1,700 frames per fibre were analysed using a modification of a previously written processing algorithm in interactive data language environment (8).

### EDL skinned fibre solutions

A single large (top 30% diameter of the fibres) intact EDL fibre was selected from the population of fibres using a fine bore pipette. We have previously shown that in mice there is a strong correlation between fibre size and type with fast fibres having nearly twice the cross-sectional area (CSA) compared to slow-twitch type 1 (9). The selected fibre was tied onto a sensitive force transducer of the *MyoRobot* biomechatronics system (19). After tying, it was placed for 10 min in solution A (see later) with 2% Triton X-100 added to chemically skin all remaining membranous cell elements. The fibre was then exposed to a series of solutions of different free Ca^2+^ concentrations. The strongly buffered Ca^2+^ solutions were prepared by mixing specific proportions of EGTA-containing solution (solution A) and Ca-EGTA– containing solution (solution B). Solution A contained 117 mM K^+^, 36 mM Na^+^, 8 mM adenosine triphosphate (ATP, total), 1 mM free Mg^2+^, 10 mM creatine phosphate, 50 mM EGTA (total), 60 mM N-[2-hydroxyethyl] piperazine-N’-[2-ethanesulfonic acid] (HEPES), and 1 mM NaN_3_ (pH 7.10). Solution B was similar to solution A, with the exception that the EGTA and Ca^2+^-EGTA concentrations of solution B were 0.3 and 49.7 mM, respectively. The free Ca^2+^ concentrations of the solutions were calculated using a K_apparent_ for EGTA of 4.78 × 10^6^ M^−1^ (20). Maximal force was determined by exposure to solution B, containing a free Ca^2+^ concentration of 3.5 × 10^−5^ M. Force was returned to baseline after maximal activation by exposure to solution A. The plateaus of the force responses elicited by exposure to solutions of increasing free Ca^2+^ concentration are expressed as a percentage of maximum Ca^2+^-activated force and plotted as a function of pCa. The force–pCa data were fitted with Hill curves using GraphPad Prism8.

### The MyoRobot, automated biomechatronics system

For full details of the *MyoRobot* see Haug (19). The following procedures were carried out on the EDL fibres using the *MyoRobot*. <u>*Force-pCa*</u>: the fibre was immersed in wells containing highly-EGTA buffered internal solutions with decreasing pCa values, made up by mixing solutions A&B (see above). Exposure to each pCa was for 20 seconds.

#### Slack test; speed of shortening

The slack test assumes a constant shortening velocity of muscle fibres upon imposing a sudden small slack to the fibre when isometrically activated. Fibres were held at resting length L_0_, transferred to a maximally activating Ca^2+^ solution and maintained in this solution until the force produced by the fibre reached a steady-state plateau. The voice coil actuator (with fibre attached) was then linearly moved at maximum speed (250 mm/s) towards the transducer pin (other end of fibre attached) for a given slack length (5–40% L_0_). While force declined to zero, the force was continuously monitored at 2 kHz high sampling rate until force redeveloped through ongoing fibre shortening, re-establishing isometric force production. When the next steady-state force level was reached, the preparation was dipped in high EGTA relaxing solution where the voice coil pin was returned to L_0_ under relaxing conditions before the next slack test was imposed.

#### Passive axial elasticity, resting length-tension curves

To assess axial fibre compliance through resting length-tension curves when the fibre was relaxed in low Ca^2+^ the voice coil was driven at very slow speed (quasi-static) to stretch the fibre while passive restoration force was sampled at 200 Hz. Since the skinned fibres possess viscous properties (e.g., presence of titin), the stretch velocity was optimized to values slow enough to be in a steady-state between instantaneous elastic restoration force and viscous relaxation.

#### Eccentric contractions

The fibre was placed in a maximal Ca^2+^ activating solution and allowed to produce maximal isometric force; it was then stretched by 20% of L_0_ for two seconds, the stretch was released for a further two seconds before the fibre was relaxed in a low Ca^2+^, high EGTA solution. The procedure was carried out three times in total, and a final maximal Ca^2+^-activating force recorded.

#### Second harmonic generation imaging of single fibres

Single EDL fibres were tied to thin glass rods and fixed in 0.1% glutaraldehyde solution for SHG microscopy. Glass rods with one EDL fibre each were mounted into a microscopy chamber immobilized between Vaseline® stripes for multiphoton imaging, for details see Friedrich (21).

### Statistics

Data were presented as means ± SD. Differences occurring between genotypes were assessed by one-way ANOVA with respect to genotype. Post hoc analysis was performed using Holm-Sidak’s multiple comparisons test. The Logrank test was used to compare survival distributions of muscle fibres during contraction and the Mann Whitney test used for comparing angular variability of myofibres between groups. All tests were conducted at a significance level of 5%. All statistical tests and curve fitting were performed using a statistical software package Prism Version 8 (GraphPad, USA).

## Results

FDB muscles were dispersed into intact single fibres by collagenase digestion. A typical digest normally yields over 200 viable fibres. Supplementary figure A shows the range of muscle fibre widths obtained from a digest from a control FDB muscle. Since we first described using Bekoff and Betz (22) digest technique on mouse fibres in 1990 (23) it has proved a robust tool to generate intact isolated mouse muscle fibres for the study of their cell physiology (24, 25). A viewing of the video from our high-speed camera of a single unloaded FDB fibre contracting at 20 Hz shows the reliability of this preparation in being able to produce repetitive unloaded contraction and relaxation cycles (Supplementary Figure B (video)). Figure 1A, B shows single FDB fibres from *Actn3KO* and WT being stimulated at 10-100 Hz and their associated fibre length and shortening velocity. The combined data is shown in Figure 1C, D. The shortening length and maximum velocity of shortening were not significantly different between *Actn3KO* and WT, however, at higher frequencies of stimulation (20-100Hz) there was a significant slowing of the minimum relative shortening length, which was similar for both genotypes Figure 1C. At 30 Hz the absolute maximum velocity was significantly faster in *Actn3KO*, this difference was no longer present at 100Hz, Figure 1D. The values we measured for velocity of shortening were similar to those previously reported for mouse fast-twitch fibres (8). For the skinned fibre experiments, we used the EDL fast-twitch muscle with longer fibres suitable for tying to a sensitive force transducer (not feasible for the ~500 μm long FDB fibres). The mouse EDL has been shown to have a fibre type distribution that is ~79% type 2B (fast glycolytic), ~16% type 2X and ~4% type 2A (fast oxidative glycolytic) muscle fibres (26).

**Figure 1:**
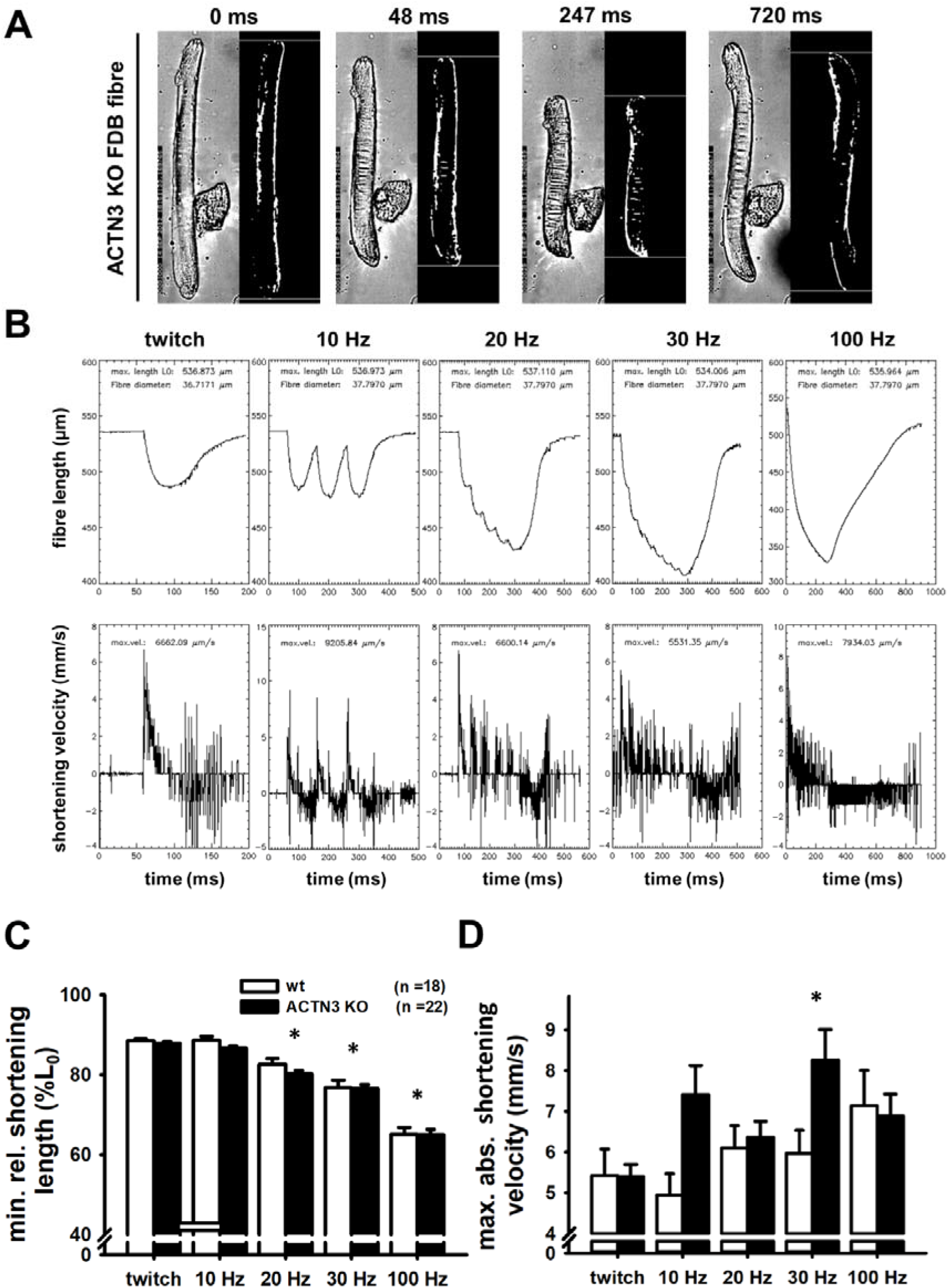
Unloaded speed of shortening of single dissociated intact FDB fibres from WT and *Actn3KO* mice. **(A),** Example images recorded with a high-speed camera during a shortening sequence of a *Actn3KO* single fibre stimulated at 100Hz and recorded at 4,166 fps at indicated time stamps. Also shown are the automated analysis images from which the shortening parameters, fibre length and shortening velocity were obtained. (**B),** Analysed time traces of these parameters for the same fibre at indicated stimulation frequencies. **(C)** Minimum shortening length was not different between genotypes, however, there was a significant overall reduction 20-100Hz as indicated *. **(D)** Absolute maximum shortening velocities were not different between genotypes apart from 30 Hz where *Actn3KO* were significantly faster as indicated by *. Data are from four animals each. Significance WT vs. KO based on one-way ANOVA test indicated as follows: * p< 0.05

Figure 2 shows the maximal force produced by isolated myofibres from *Actn3KO* and WT mouse EDL. Figure 2A shows that that *Actn3KO* fibres tended to produce less absolute force than WT, but when corrected for CSA (specific force) the maximal force output of the myofibrillar proteins was the same for both genotypes, confirming our results from the intact whole EDL (15). Figure 2B shows the fibre diameter distributions of WT and *Actn3KO*; here we see a trend for the fibres to be of smaller diameter in *Actn3KO* as we have previously reported (9), however, here these are not random samples, as for both WT and *Actn3KO*, we actively selected the longest fibres with the largest diameters for attaching to the force transducer. Figure 2C shows the combined pCa-force curves generated for the WT and *Actn3KO* fibres; there were no meaningful differences in the contractile properties, i.e. the slope of the pCa-force curves or pCa_50_ and these parameters were in the range of those previously reported for fast-twitch fibres (27).

**Figure 2:**
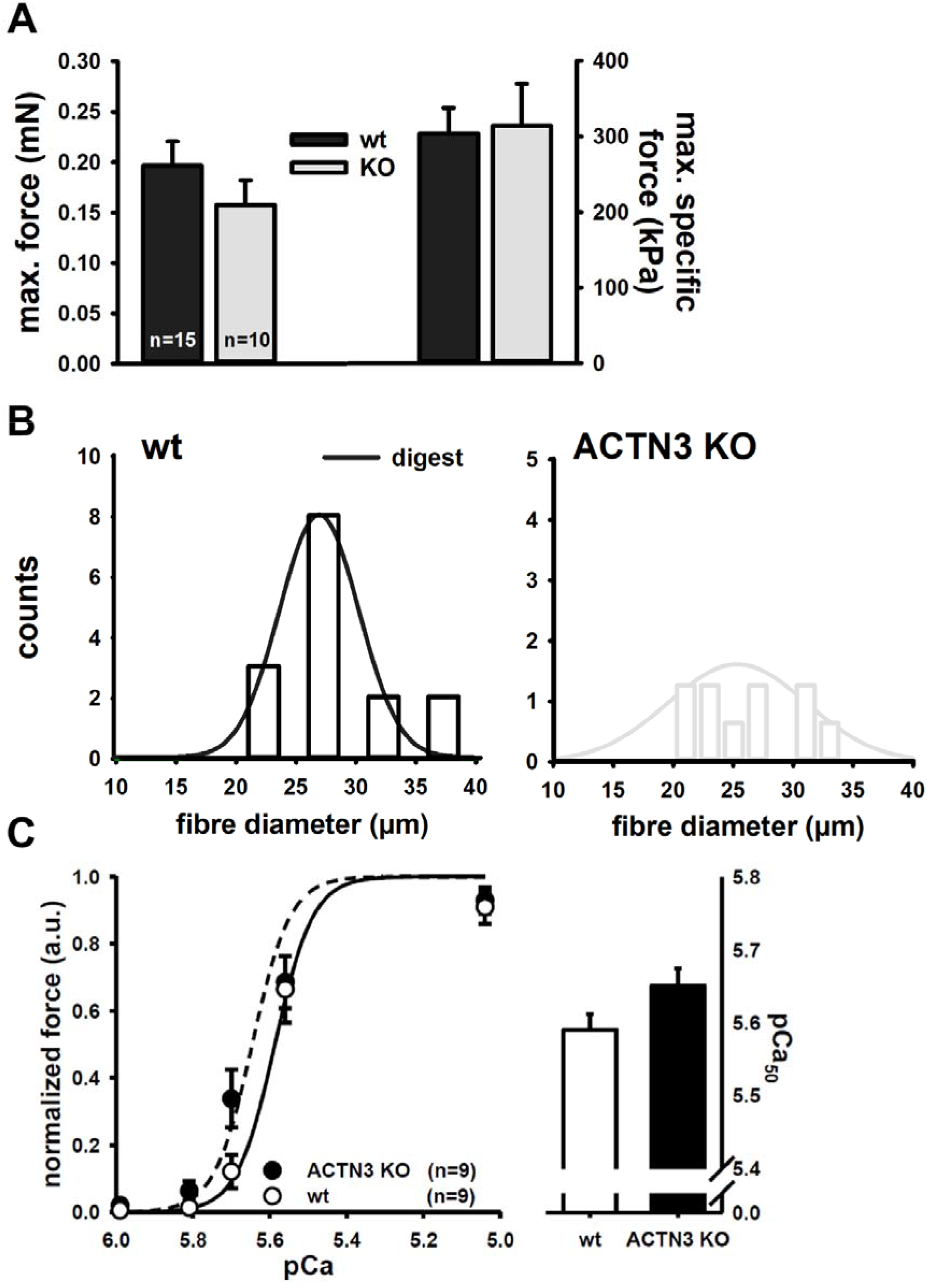
Maximum myofibrillar force in single EDL fibres from *Actn3KO* mice is unaltered compared to WT. **(A),** Statistical analysis of maximum force and specific force (normalized to fibre diameter-derived CSA) values from EDL muscles of WT and *Actn3KO* mice. No significant differences based on one way ANOVA tests were apparent. (**B),** Analysis of fibre diameter distributions shows a trend to smaller diameter in the *Actn3KO* fibres. (**C)**, Calcium-sensitivity shown as pCa-force relationship. Average data are displayed along with the reconstructed average fit, which suggest a shift towards a decreased Ca^2+^-sensitivity in *Actn3KO* fibres.

In Figure 3, we show the results for unloaded velocity of shortening (19) where a fibre is first activated in the maximally activating pCa solution and then subject to a series of rapid releases of tension at release ‘slack’ lengths 10-40% of L_0,_ the rate of tension redevelopment is measured for each ‘slack’. We focused on the fast-phase of tension redevelopment which is related to the myosin crossbridge kinetics. Figure 3A shows the raw data run from a WT EDL fibre, the dotted line labelled F_sig_ delimits the end of the fast phase of force redevelopment resulting from the cycling of the myosin heads. In Figure 3B, we report the rate of change of length during the fast phase, the slope of the lines gives us the parameter V_fast_ (fast velocity) which is shown as a bar graph in Figure 3C demonstrating that the *Actn3KO* genotype does not alter the fast phase of the velocity of shortening.

**Figure 3:**
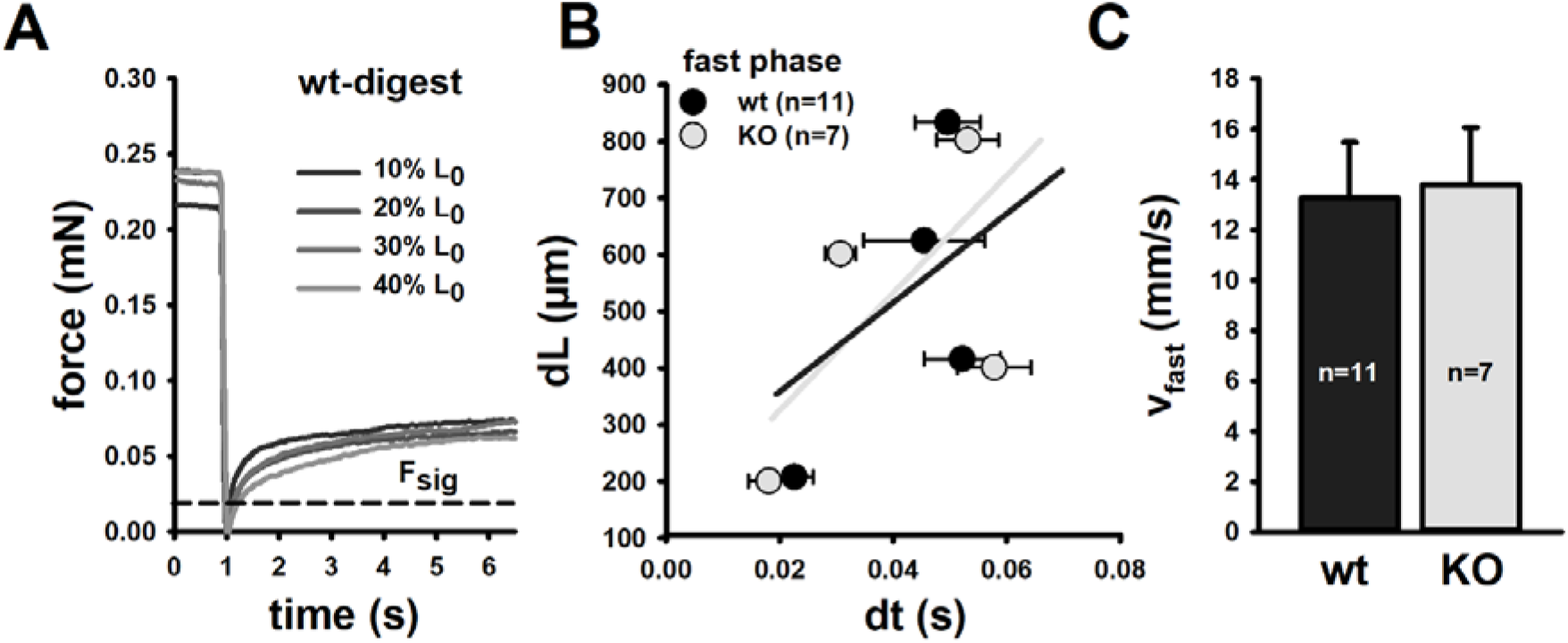
Unloaded shortening of single EDL fibres following myofibrillar activation is unaltered in *Actn3KO* but internally loaded shortening is slowed in digested single *Actn3KO* fibres. **(A),** Representative example recording of a single muscle fibre during the slack-test using the *MyoRobot* to sequentially apply a series of slacks between 10% and 40% of resting length L_0._ **(B)**, dL-dt relationships of 7 and 11 single fibres from WT and *Actn3KO* mice reveals similar shortening kinetics in both. (**C),** Statistical analysis based on a one-way ANOVA of V_fast_ values supports the findings of similar shortening kinetics for the entire data set.

We further demonstrate that resting length-tension curves and steady-state compliance are not different between *Actn3KO* and WT myofibres, showing that the absence of α-actinin-3 from the Z-discs did not alter the elastic properties of the myofibrils (Figure 4C.).

**Figure 4:**
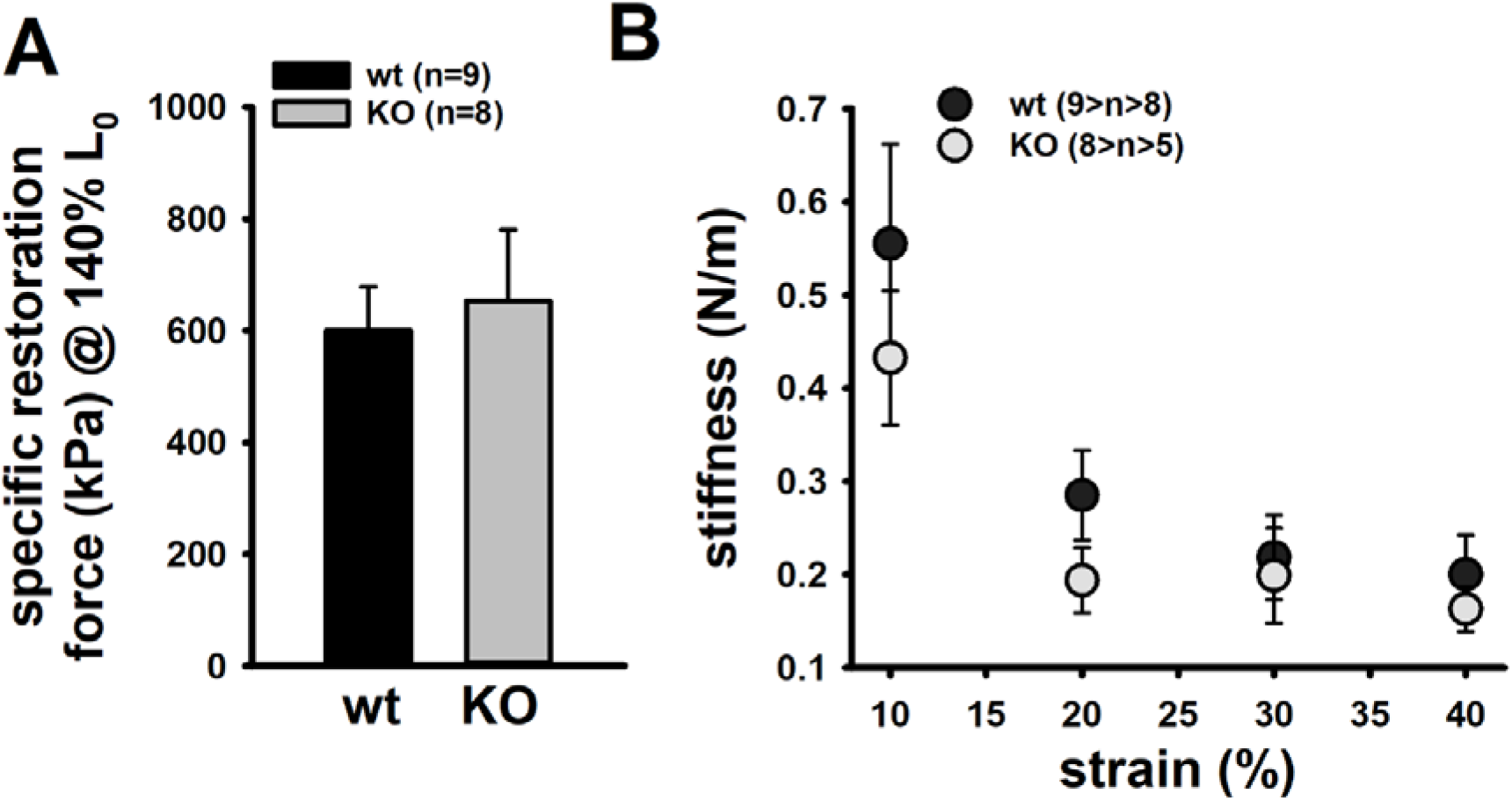
Resting length-tension curves and steady-state compliance of EDL *Actn3KO* single fibres. **(A),** Specific restoration force (stress) at 140% L_0_ analysed in a number of WT and *Actn3KO* fibres. (**B),** Steady-state stiffness values vs. strain indicate somewhat lower mechanical stiffness in the *Actn3KO* background compared to the WT.

We next looked at the single fibre visco-elasticity in *Actn3KO* and WT by rapidly stretching the single fibre in a series of 10% steps up to 60% longer than its starting value of L_0_ (100%) (Figure 5). Here, a peak restoration tension was rapidly reached with each step followed by an exponential force relaxation as shown in Figure 5A. Three parameters; absolute specific restoration force, absolute specific relaxation force and the rate of relaxation are shown on the raw data trace in Figure 5A. Total maximal specific force produced by each stretch was not different between *Actn3KO* and WT (Figure 5B). After attaining maximum force at each length, the fibre was allowed to relax to a new steady-state over four seconds. Then, we evaluated the amount of force drop during the relaxation period, which was not significantly different between WT and *Actn3KO* (Figure 5C). Figure 5D shows the rate of relaxation, and once again, there was no difference between WT and *Actn3KO*.

**Figure 5:**
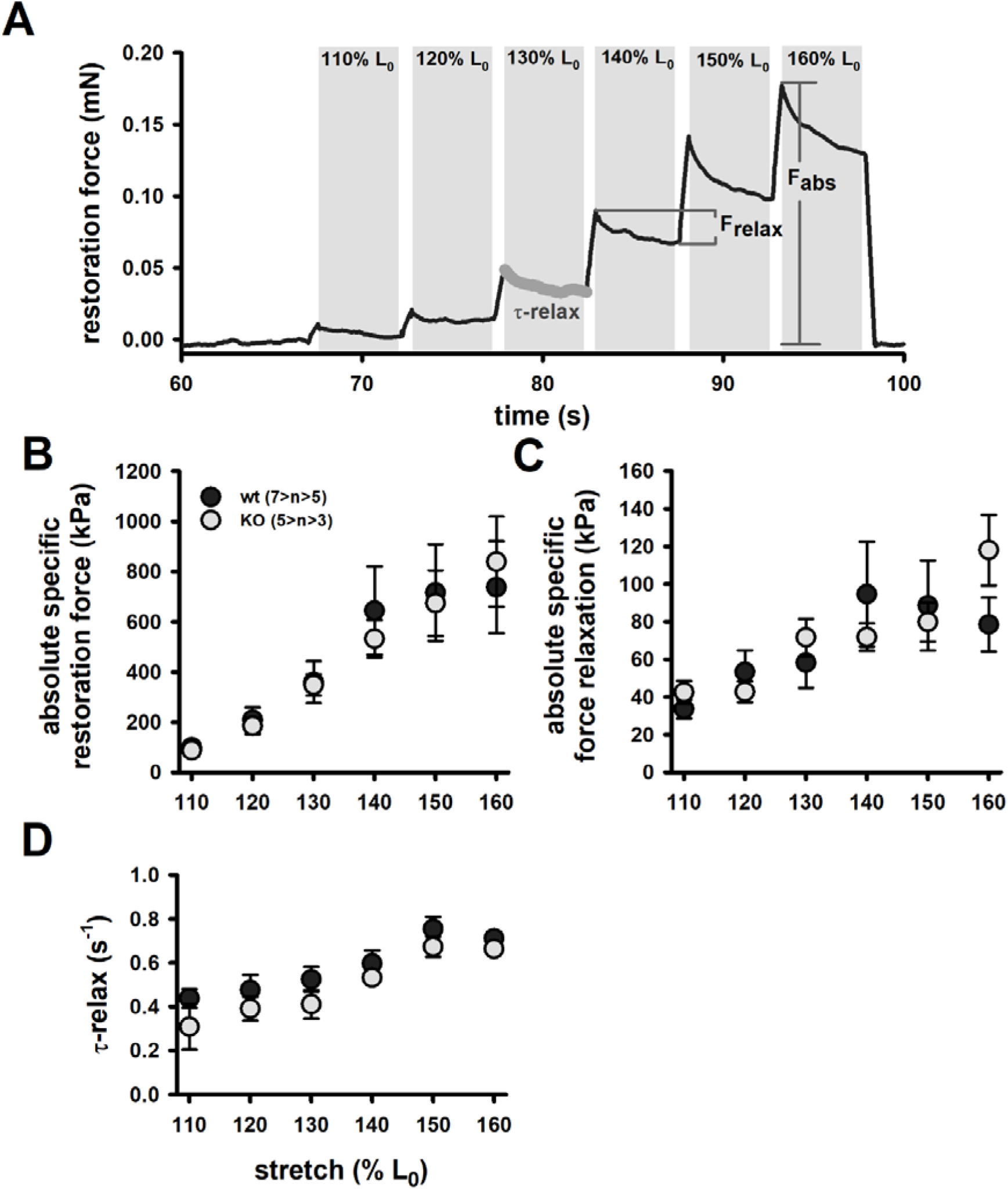
Single fibre visco-elasticity in EDL muscle from adult and old *Actn3KO* mice. **(A),** Example trace of ‘strain-jumps’ of increasing 10% L_0_ ampitudes quickly applied to the fibre using the *MyoRobot* biomechatronics system. Each sudden stretch is answered by an instantaneous increase in restoration force F_R_ to a new maximum F_abs_ **(B)** before exponentially relaxing to achieve a new steady-state level with relaxation amplitude F_relax_ **(C)** and a time constant τ_relax_ **(D)**.

To investigate the mechanical stability of the *Actn3KO* fibres, a set of three eccentric contractions were performed at +20% of L_0_ (Figure 6A), the fibre was first maximally activated by exposing it to a high Ca^2+^ solution. Once the force had plateaued it was stretched by 20% of L_0_, held for two seconds and then released. The fibre was then allowed to reach a new maximal plateau for two seconds before being relaxed in a high EGTA relaxing solution. Figure 6B shows that during the three contraction eccentric protocol, there was a significant number of fibres which broke apart so that the fibre separated into two distinct pieces. When we quantified these breakages, it was clear *Actn3KO* fibres broke more frequently because of the eccentric contraction protocol than WT (Figure 6B).

**Figure 6:**
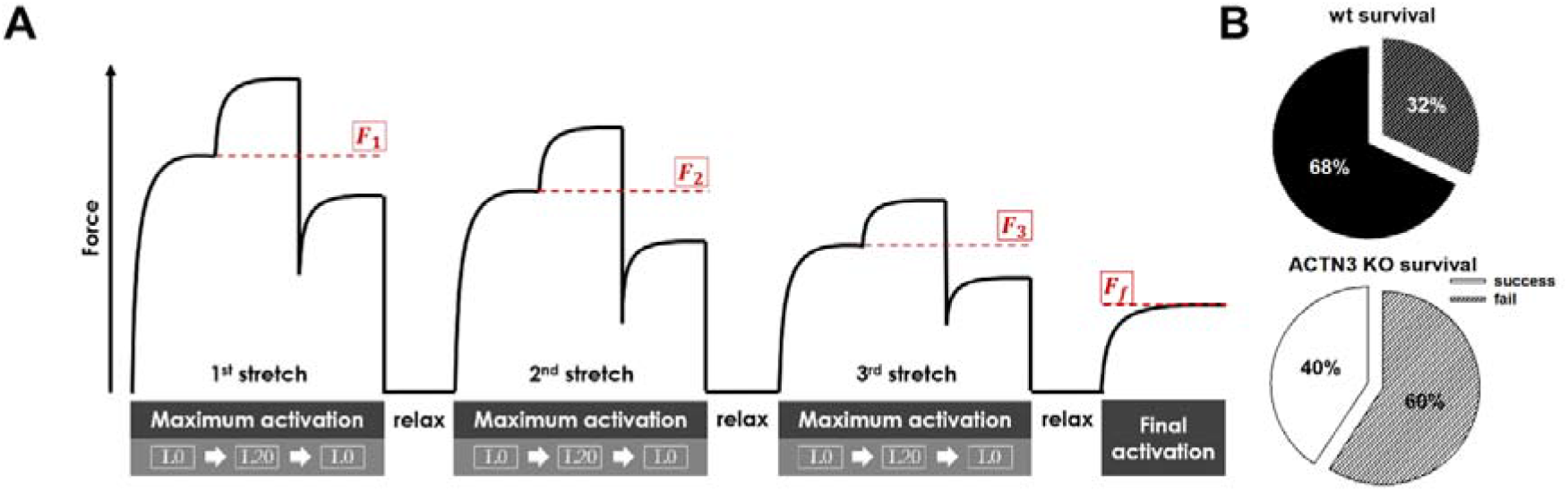
Contraction induced breakages are more pronounced in EDL fibres from *Actn3KO* mice. **(A),** Scheme of recurrent eccentric contractions, maximally activating a single fibre and then imposing a 20% stretch for several seconds before returning to resting length L_0_ and subsequent relaxation. Three such eccentric contractions were pursued followed by a final assessment of maximum isometric force. (**B)**, *Actn3KO* fibres showed much lower survival and higher rate of breakage during the sequence to a significance of p=0.02 based on logrank analysis.

To investigate if there was a morphological reason for the increased breakage in the *Actn3KO* fibres, we used *Second Harmonic Generation imaging* (SHG) and quantitative morphometry in single EDL muscle fibres (Figure 7). Group analysis of 54 WT fibres and 35 *Actn3KO* fibres showed that the *Actn3KO* fibres had significantly higher levels of myofibrillar axial lattice disorder, which we term vernier density (VD) likely due to Z-disc anchorage inhomogeneities resulting from the absence of α-actinin-3 (Figure 7B). These myofibrillar Y shaped vernier deviations or disruptions (Figure 7A, B) will be points susceptible to mechanical weakness in the contractile filaments present in fibres (see discussion for modelling).

**Figure 7:**
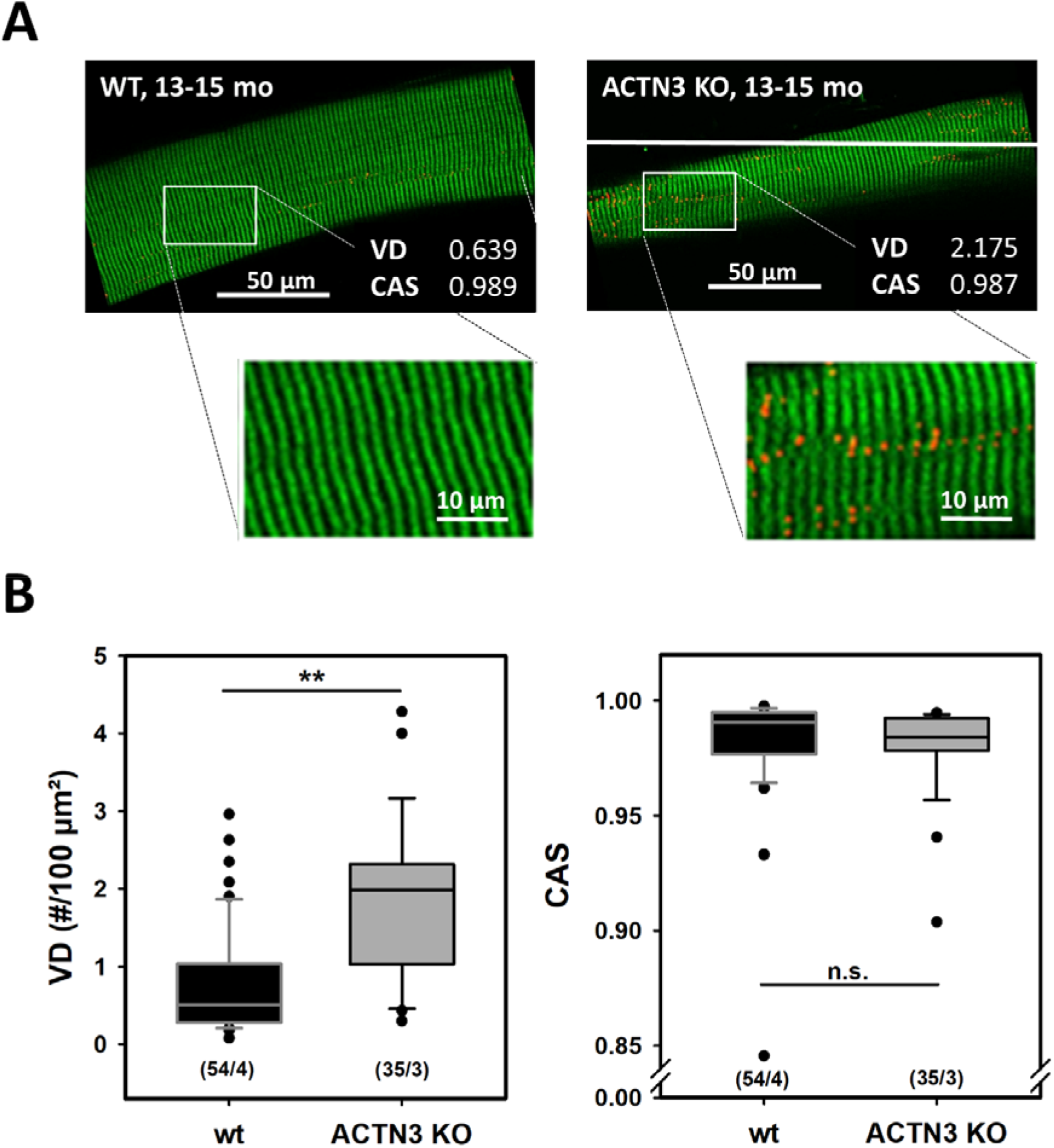
SHG imaging and quantitative morphometry in single dissociated EDL fibres reveal misregistered myofibrillar ultrastructure in *Actn3KO*. **(A),** Representative example images from the middle plane of a single WT and *Actn3KO* EDL fibre (top) of the same 13-15 month age group alongside with the automatically detected verniers (red) and the vernier density (VD) and cosine angle sum (CAS) values derived from the morphometry analysis. A magnified rectangular section is given below the images. (**B),** Group analysis in a substantial number of single fibres from several animals reveals significantly higher VD values in *Actn3KO* fibres over fibres from WT littermates, indicative of a higher linear *out-of-register* disorder. As for the angular variability of myofibrils, the CAS values were similar in both groups. **: p < 0.001, Mann-Whitney Rank sum test. (n/m): n single fibres from m animals.

## Discussion

Force transmission from the myosin heads to the Z-disks (a major component of which is the protein of α-actinin-2 and α-actinin-3) is mediated by actin filaments and titin (28). The Z-discs are the focal point of force transmission and mechanical strength within the fibre (5). When muscles are damaged by excessive forces, such as those experienced during an eccentric contraction, electron micrographs show that the initial point of damage is at the Z-disc (29). Fast-twitch muscles express their own isoform of α-actinin, α-actinin-3. Globally, ~1.6 billion people have a polymorphism (R577X) in the *ACTN3* gene which means they cannot produce the protein α-actinin-3 in their fast-twitch muscles (2); in these people, α-actinin-2 is upregulated to (partially) compensate for the loss. We have generated an *Actn3KO* mouse model to study the morphological and contractile consequences of the absence of α-actinin-3 from fast-twitch muscles. However, it should be born in mind that the Z-disc also has a key role as a force sensor linking tension along the myofibrils with intracellular chemical signalling mechanisms (30). Our FDB digest technique has been refined from our first digest of mouse skeletal muscle fibres in 1990 (23). In its current form the digest produces over 200 single contracting fibres per batch, the majority of which could undergo at least three consecutive rounds of fatiguing contractions followed by recovery. High magnification high speed video framing (up to 4,166 frames per second) showed that over 70% of the single FDB fibres attached to cleaned glass cover slips at their centre portions. They shortened and relaxed linearly about this point (Figure 1A&B and supplementary Figure B (video)) when stimulated with a supramaximal voltage pulse 1 ms in duration delivered from a bipolar pair of platinum wires insulated to the tips and positioned close to the neuromuscular junction. A portion of the fibres would bend into crescent shapes during repeated contractions, and those were discarded from shortening analyses. A platinum electrode was used to stimulate the selected fibres from 10-100 Hz, we showed the maximum shortening velocity and minimum shortening length were basically the same in *Actn3KO* fibres and WT. This supports our earlier findings which reported the *Actn3* polymorphism and resulting absence of α-actinin-3, did not alter the expression of the fast myosin isoform (9, 10, 15), however, this is the first time this has been shown directly for unloaded shortening of fibres rather than inferred from the myosin type. For the skinned fibre experiments, we used the EDL muscle to get individual fibres which were long enough to manually tie to a force transducer biomechatronics system. Given ~79% of EDL fibres are of 2B MHC isoform, ~16% type 2X and ~4% type 2A (26), we have previously shown (9) that 2B fibres are around twice the diameter of the 2X and 2A fibres. Thus, by selecting the top 30% of the largest diameter fibres, we were confident in having 2B fibres. This was confirmed by the pCa-force curves which were consistent with fast-twitch type 2B fibre types (Figure 2C). The similarity of the pCa-force curves between genotypes supports earlier findings that there are no changes in myosin isoforms (1). α-Actinin proteins are a key component of the Z-discs anchoring actin fibres from adjacent sarcomeres and transmitting force longitudinally to the tendons (11, 28), thus the chemical skinned fibre technique, using whole chemically skinned fibres (as opposed to mechanically skinned fibre segments) is the ideal way to test the effect of absence of α-actinin-3 from the Z-discs in fast-twitch muscle. In this preparation there is no interference from adjacent fibres, connective tissue or lateral transmission of force (28). We measured unloaded shortening, resting length tension, stiffness, and visco-elasticity using the *MyoRobot* biomechatronics skinned fibre set up (19).

We found no significant differences in these properties indicating that both α-actinin-2 and α-actinin-3 confer the same mechanical properties to Z discs. In an elegant, skinned fibre study on *ACTN3KO* humans, Broos (31) showed in terms of their visco-elastic properties, fast-twitch 2X muscle fibres from α-actinin-3 positive humans were the same as those from humans who were α-actinin-3 negative. The group also showed that there was no difference in maximum specific force with respect to genotype, as we would predict from our earlier studies where we showed there was no change in myosin isoform expression with respect to genotype (1, 9, 10, 15). Our current studies in the mouse confirm the human findings, and while we did see a trend towards less absolute force in *Actn3KO*, this was due to the fact, as confirmed by Broos (31), that *Actn3KO* fast fibres (2X humans, 2B mice) are significantly smaller in diameter, and when we normalised the force results for cross sectional area, we found the maximal specific force was the same.

We have previously reported that when the isolated EDL muscles are subjected to five eccentric contractions of 20%+L_0_ strain there is no effect of genotype on the eccentric contraction-induced force deficit (15). Intriguingly, in a later study (16) when we used a stronger eccentric contraction protocol with a strain of 30%+L_0,_ there was a significant increase in the eccentric contraction force-deficit in the *Actn3KO* fast-twitch EDL muscles. The results from the current study provide a likely explanation for these whole muscle results as we show that in some cases fibres from both *Actn3KO* and WT EDL muscles can withstand three eccentric contractions of 20% strain, however, ~60% of the *Actn3KO* fibres broke during the procedure compared with ~35% of the WT fibres (Figure 6B). *Second harmonic imaging* of single EDL fibres revealed a myofibrillar structural deformity which provides a plausible explanation for this increased mechanical instability associated with the *Actn3KO* genotype. *Actn3KO* fibres contained numerous axial lattice shift of adjacent myofibrils (verniers apparent as ‘Y’-patterns) of the myofibrillar contractile proteins, like the vernier deviations we have previously reported in unbranched dystrophic muscle fibres (termed “chaotic”) from the *mdx* mouse (21, 32). Modelling from intact fibres (33) has shown that macroscopic splits or branches within a fibre, are points of mechanical weakness which may be susceptible to breakage when fibres were stressed by eccentric contractions. We propose that the *Actn3KO* fast fibres behave normally under moderate strains, but as the strain increases there comes a point where the weaker dislodged myofibril arrays start to snap, setting up a positive feedback loop placing additional stresses on the remaining VD which in turn break. This provides an explanation as to why the intact muscle was not damaged at 20%+L_0_ eccentric contraction strain (15), but showed a significant force loss at 30%+L_0_ (16), which in our model would be sufficient strain to rupture the weaker myofibrillar out-of-register lattice in the fast α-actinin-3 deficient fibres. We have previously shown the presence of internalized centralized nuclei at baseline in *Actn3KO* muscles. Centralized nuclei were not present in the age matched controls (16). Centralized nuclei are an accepted histological marker of a regenerated fibre. This would suggest these fast-twitch fibres with increased myofibrillar lattice shifts are subject to damage during normal muscle contraction when compared to WT fast fibres containing α-actinin-3 in the Z-discs. There have been several reports in the literature that α-actinin-3 deficient individuals may experience faster decline in muscle function with increasing age (9, 34) and our results may explain some of this decline because as sarcopenia develops, the loss of muscle mass will place greater stress on the remaining fast-twitch muscles during eccentric contractions. In the case of individuals lacking α-actinin-3 protein, their remaining fast-twitch muscles will be at greater risk of damage compared to α-actinin-3 protein positive individuals. This will be compounded by the reduced diameter of the fast-twitch muscles in the α-actinin-3 protein deficient individuals, increasing mechanical axial stress on single fibres.

## Conclusion

1. Unloaded single fibre shortening length and maximum speed of shortening at different field-stimulation frequencies (10-100Hz) are unaltered by the absence of α-actinin-3 in *Actn3KO* single intact muscle fibres from FDB fast-twitch muscle.
2. Using the chemically skinned fibre technique with single fast-twitch EDL fibres, we show visco-elastic properties and myofibrillar force production (force-pCa) are not affected by the absence of α-actinin-3.
3. When chemically skinned single EDL fibres were maximally activated and subjected to three eccentric contractions of Lo+20% strain, ~60% of the *Actn3KO* fibres broke during the procedure compared with ~35% of the WT fibres.
4. Second harmonic imaging of single *Actn3KO* EDL fibres revealed myofibrillar structural abnormalities with an axial lattice shift of adjacent myofibrils (verniers apparent as ‘Y’-patterns).
5. The structural weakness caused by the Y shaped vernier branches provides a plausible explanation for the increased mechanical instability associated with the *ACTN3* genotype.

## Additional files

**Additional file 1: Supplementary figure A.** Frequency distribution of FDB fibre widths. A total of 200 WT FDB fibres were measured.

**Additional file 2: Supplementary figure B.** High-speed camera view of a single unloaded FDB fibre contracting at 20 Hz.

## List of abbreviations

CAS: Cosine angle sum
CSA: Cross sectional area
dL: Distance shortened
dT: Time shortened
EDL: Extensor digitorum longus
F_abs_: Maximum absolute force
F_R_: Restoration force
F_relax_: Force at steady state
F_sig_: End of fast phase
FDB: Flexor digitorum brevis
K_apparent_: Apparent dissociation constant
KO: Knock out
L_0_: Optimal length
MyHC: Myosin heavy chain
pCa: Calcium concentration
pCa_50_: Calcium concentration at 50% of maximum force
SHG: Second harmonic generation
τ_relax_: Time to reach steady state
VD: Vernier density
V_fast_: Fast velocity
WT: Wild type

## Declarations

### Ethics approval and consent to participate

Use of animals was approved by the Animal Care and Ethics Committees of the Children’s Medical Research Institute and the University of New South Wales.

### Consent for publication

Not applicable, no data was obtained from individual person

### Availability of data and materials

The datasets used and/or analysed during the current study are available from the corresponding author on reasonable request.

### Author’s contributions

MH, BR, SN, LK, DAGM, PJH, KNN, OF and SIH Conceived of and designed research;

MH, BR, SN, LK, DAGM, OF and SIH Performed the research;

MH, BR, SN, LK and DAGM Analysed the research;

SIH Interpreted results of experiments and drafted the manuscript;

MH and OF Prepared figures;

MH, BR, LK, DAGM, PJH, KNN, OF and SIH Edited and revised the manuscript.

All authors approved the final version of manuscript and agree to be accountable for all aspects of the work in ensuring that questions related to the accuracy or integrity of any part of the work are appropriately investigated and resolved. All persons designated as authors quality for authorship, and all those who qualify for authorship are listed.

