## Supplementary figures and images for "Absence of the Z-disc protein α-actinin-3 impairs the mechanical stability of *Actn3KO* mouse fast-twitch muscle fibres without altering their contractile properties or twitch kinetics"

### Supplementary A

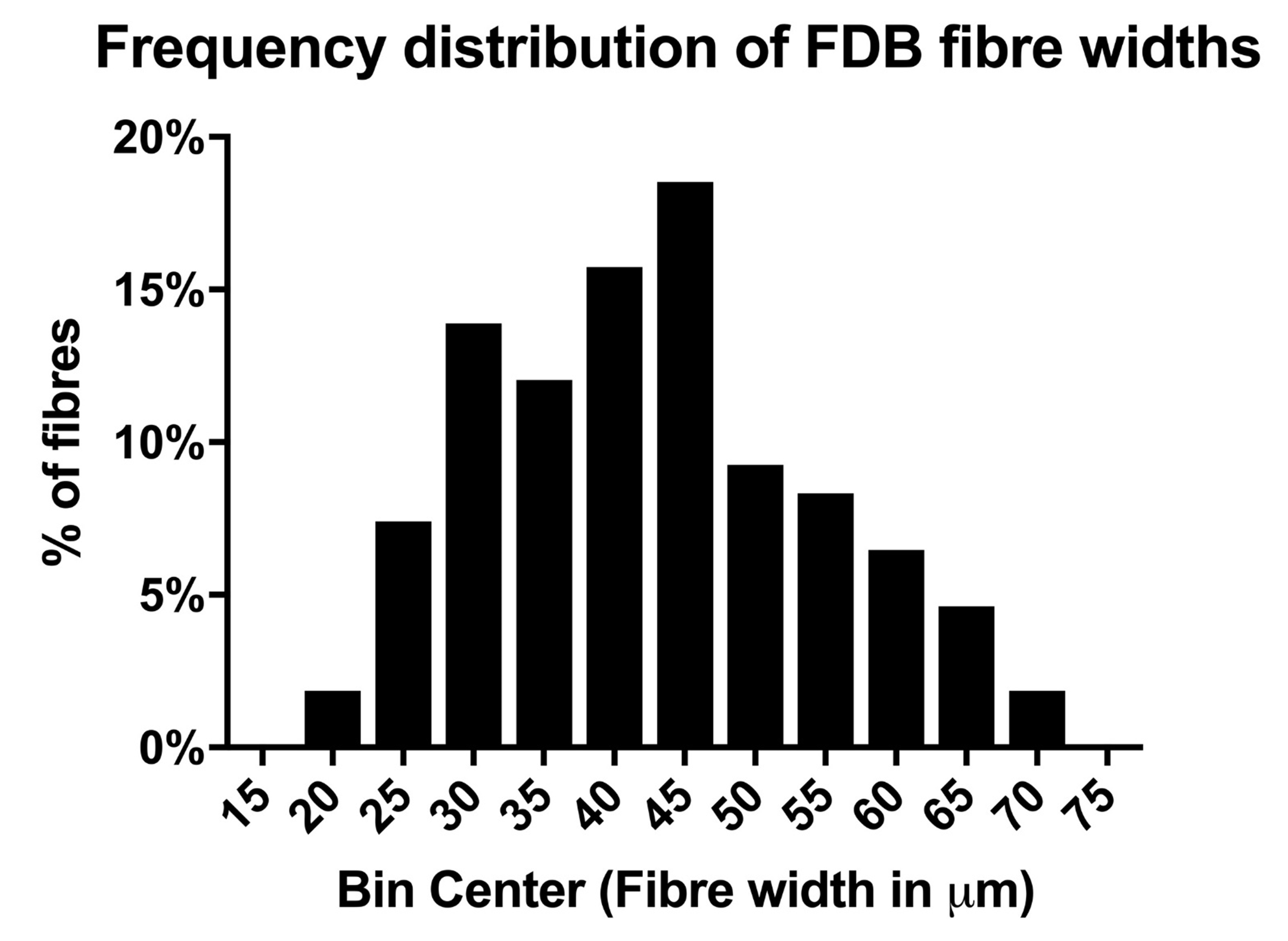
